# Structural analysis of the OC43 coronavirus 2′-O-RNA methyltransferase

**DOI:** 10.1101/2021.03.16.435745

**Authors:** Pavel Dostalik, Petra Krafcikova, Jan Silhan, Evzen Boura

## Abstract

The OC43 coronavirus is a human pathogen that usually causes only the common cold. One of its key enzymes, similar to other coronaviruses, is the 2′-O-RNA methyltransferase (MTase) that is essential for viral RNA stability and expression. Here, we report the crystal structure of the 2′-O-RNA MTase in a complex with the pan-methyltransferase inhibitor sinefungin solved at 2.2 Å resolution. The structure revealed an overall fold consistent with the fold observed in other coronaviral MTases. The major differences are in the conformation of the C-terminus of the nsp16 subunit and an additional helix in the N-terminus of the nsp10 subunits. The structural analysis also revealed very high conservation of the SAM binding pocket suggesting that the SAM pocket is a suitable spot for the design of antivirals effective against all human coronaviruses.

**Importance:** Some coronaviruses are dangerous pathogens while some cause only common colds. The reasons are not understood although the spike proteins probably play an important role. However, to understand the coronaviral biology in sufficient detail we need to compare the key enzymes from different coronaviruses. We solved the crystal structure of 2′-O-RNA methyltransferase of the OC43 coronavirus, a virus that usually causes mild colds. The structure revealed some differences in the overall fold but also revealed that the SAM binding site is conserved suggesting that development of antivirals against multiple coronaviruses is feasible.

## Introduction

Prior to the COVID-19 pandemic only six other human coronaviruses were known - the already extinct deadly SARS-CoV, MERS-CoV, and four other coronaviruses (OC43, 229E, NL63 and HKU1) that are responsible for about 30% of mild respiratory diseases [1]. All of these are members of the subfamily *Orthocoronavirinae* of the *Coronaviridae* family. The *orthocoronavirinae* subfamily is further divided into four genera: *Alpha-, Beta-, Gamma-,* and *Deltacoronavirus*. The 229E-CoV and the NL63-CoV are members of the genus *Alphacoronavirus* while the OC43-CoV, HKU1-CoV, SARS-CoV, SARS-CoV-2, and MERS-CoV are all members of the genus *Betacoronavirus* [2].

The OC43 (Organ Culture 43) coronavirus has a 30.7 kbp long positive-sense single stranded RNA (+RNA) genome. That is unusual for a +RNA virus but similar to other coronaviruses [3]. It was transmitted recently, in the 19th century, to humans probably from cattle; the bovine CoV is its closet relative. It has also been speculated that OC43-CoV might be responsible for the 1889-1890 pandemic [4] which has usually been attributed to the H2N2 influenza strain [5]. Interestingly, while the OC43-CoV usually infects the upper respiratory tract and causes respiratory diseases, it is also neurotropic and can be neuroinvasive [6,7].

The coronaviral genome usually encodes for two polyproteins (pp1a and pp1ab), the spike (S), envelope (E), membrane (M) and nucleocapsid (N) and several accessory proteins. OC43, in addition, encodes for hemagglutinin-esterase (HE) that dramatically increases the infectivity of OC43-CoV [8]. The pp1a and pp1ab polyproteins are autocatalytically cleaved into 16 non-structural proteins (nsp1 - nsp16) that ensure many enzymatic activities needed for viral replication, most notably the RNA replication machinery [9]. Nsp7, nsp8 and nsp12 form the viral RNA-dependent RNA-polymerase (RdRp), nsp13 functions as a helicase and nsp10, nsp14 and nsp16 are RNA methyltransferases (MTase). Nsp14 is responsible for proofreading during RNA synthesis; it also has an additional enzymatic activity - as an exoribonuclease [10].

In this study we focused on the 2’-O MTase of OC43-CoV. RNA methylation is important for (viral) RNA stability, it protects the RNA from degradation (innate immunity shielding) and facilitates its efficient translation [11–13]. Installation of the 5’ cap in coronavirus infected cells is a four step process: 1) the 5’ gamma phosphate of the nascent RNA is hydrolyzed probably by the coronaviral nsp13 helicase; 2) GMP is transferred to the 5’ end by an unknown guanylyltransferase; 3) the nsp10:nsp14 protein complex methylates the N7 position of the newly attached guanosine; and 4) the nsp10:nsp16 protein complex methylates the 2’-O of the first nucleotide ribose. The goal is to create viral RNA that is stable in human cells, is efficiently translated and does not induce the innate immun response. Non-methylated RNA is recognized by the RIG-I (retinoic acid-inducible gene I) pattern recognition receptor and also recognized and bound to IFN-induced proteins with tetratricopeptide repeats 1 and 5 (IFIT 1 and IFIT5), which efficiently prevent its translation [14,15]. These facts imply that inhibiting the nsp10:nsp16 MTase is a valid strategy to fight coronaviral infections.

Here, we present the crystal structure of the OC43-CoV 2’-O MTase (nsp10:nsp16 complex) to understand how different coronaviruses evade innate immunity. The structure revealed relatively conserved architecture of the active site between OC43-CoV and SARS-CoV-2 and suggested that design of MTase inhibitors targeting multiple human coronaviruses is feasible.

## Results

### Crystallization of the OC43-CoV nsp10:nsp16 MTase

We analyzed the sequence of all human coronaviral nsp10 and nsp16 proteins (Figure 1). The OC43 nsp10 has 53% sequence identity and 68% sequence similarity to SARS-CoV-2, similarly the OC43 nsp16 displays 66% sequence identity and 76% sequence similarity, which is not much for viruses of the same genus. We aimed to solve the crystal structure of OC43 to the nsp10:nsp16 MTase to find out the degree of 3D structure conservation of an essential enzyme between these two coronaviruses. We used full length nsp16 and slightly truncated nsp10 (residues 10 - 131) for crystallization trials. Both proteins were expressed recombinantly and subjected to the crystallization trials. Eventually, we obtained crystals that diffracted to 2.2 Å. We solved the structure by molecular replacement and refined it to good R factors and geometry (details in Table 1 and in the M&M section).

**Figure 1.**
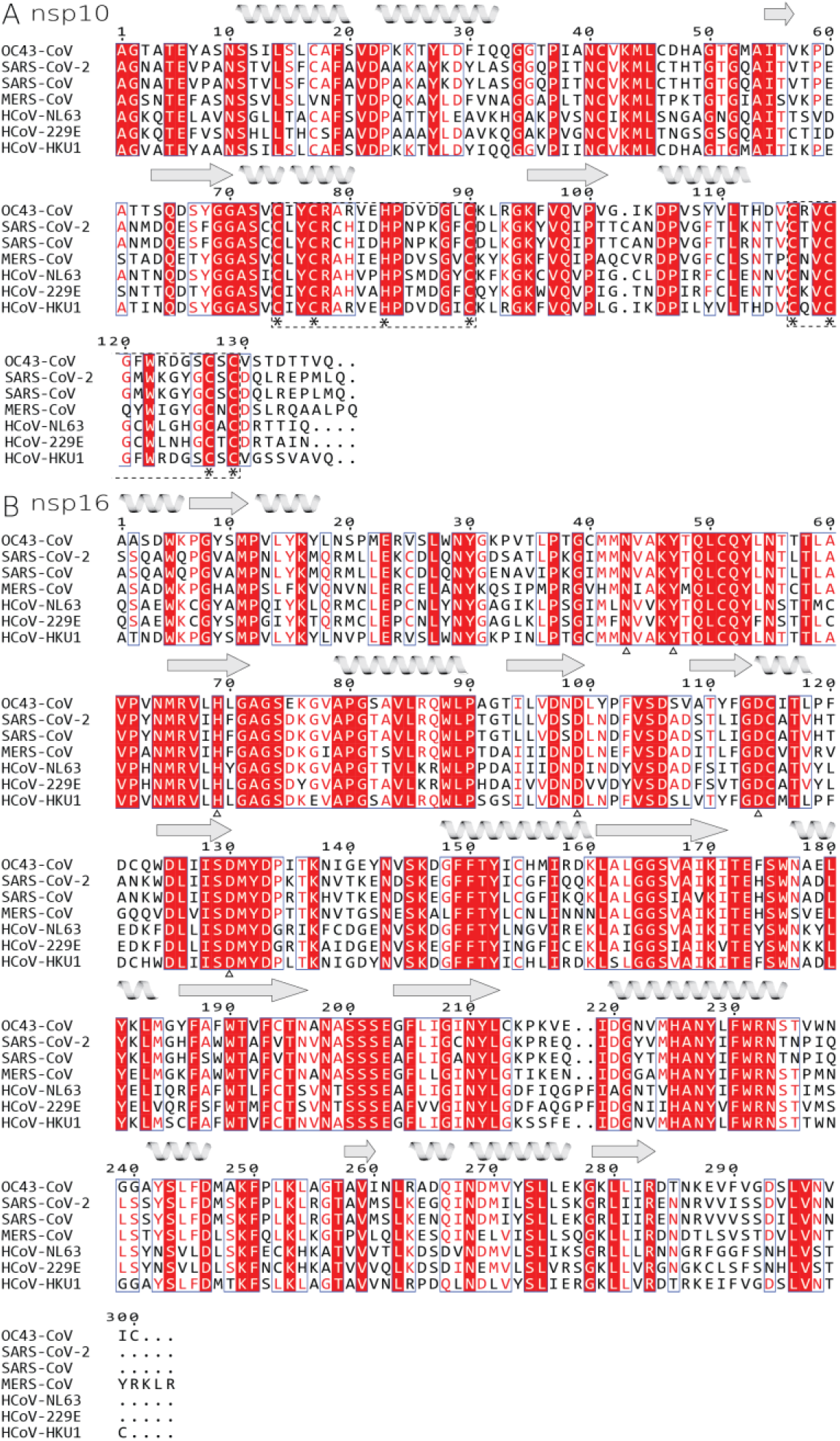
Sequence alignment of all human coronaviral nsp10 and nsp16 proteins. A) nsp10 and B) nsp16. Identical residues are highlighted in red field while conserved residues are red font. Important residues involved in coordination of zinc are in marked by asterisk below and whole zinc-binding region is in dashed box, whilst residues coordinating sinefungin are marked by empty triangle. Secondary structure features of OC43-CoV are symbolized by helices and arrows for beta sheets.

**Table 1.** Crystallographic data collection and refinement statistics. Numbers in parentheses refer to the highest resolution shell.

|  |  |
| --- | --- |
| **Crystal** | **OC43 nsp10-nsp16** |
| PDB accession code | 7NH7 |
| **Data collection and processing** |  |
| Space group | P3<sub>1</sub> |
| Cell dimensions - a, b, c (Å) | 61.9, 61.9, 109.7 |
| X-ray source | Home source |
| Wavelength, (Å) | 1.5419 |
| Resolution range (Å) | 36.57 - 2.2 (2.28 - 2.2) |
| No. of unique reflections | 23828 (2385) |
| Completeness (%) | 99.74 (99.13) |
| Multiplicity | 5.1 (4.7) |
| Mean I/$\sigma$(I) | 11.66 (2.48) |
| CC<sub>1/2</sub> | 0.981 (0.687) |
| CC* | 0.995 (0.902) |
| R<sub>merge</sub>, % | 13.05 (62.57) |
| **Structure solution and refinement** |  |
| R-work (%) | 16.41 (21.51) |
| R-free (%) | 21.01 (29.29) |
| R.m.s.d. - bonds (Å) / angles (°) | 0.005 / 0.86 |
| Average B factors (Å<sup>2</sup>) | 28.24 |
| Clashscore | 7.84 |
| Ramachandran favored/outliers (%) | 97.32 / 0 |

### The overall fold of OC43-CoV nsp10:nsp16 MTase

The overall fold is consistent with the fold of previously analyzed coronaviral nsp10:nsp16 MTases from SARS-CoV-2, SARS-CoV and MERS-CoV [16–19]. We could trace the entire protein chain except for the very last five residues of nsp16 and the last residue of nsp10. The nsp16 subunit is composed of 10 β-sheets and 11 helices (Figure 2). It exhibits a Rossmann fold where the β-sheets are arranged in a central β-motif in the shape of a letter *“J”* that is surrounded by α-helices (Figure 2B). The small nsp10 subunit is composed only of three small β-sheets and five α-helices (Figure 2B). The nsp10 fold is stabilized by two zinc fingers. The first zinc finger stabilizes the central part of the nsp10 molecule and is formed by Cys74, Cys77, His83, and Cys90. The second zinc finger stabilizes the very C-terminus and is formed by Cys166, Cys119, Cys127 and Cys129 (Figure 2D). Notably, all the residues forming both zinc fingers are absolutely conserved among human coronaviruses highlighting their importance for the function of the nsp10 protein.

**Figure 2.**
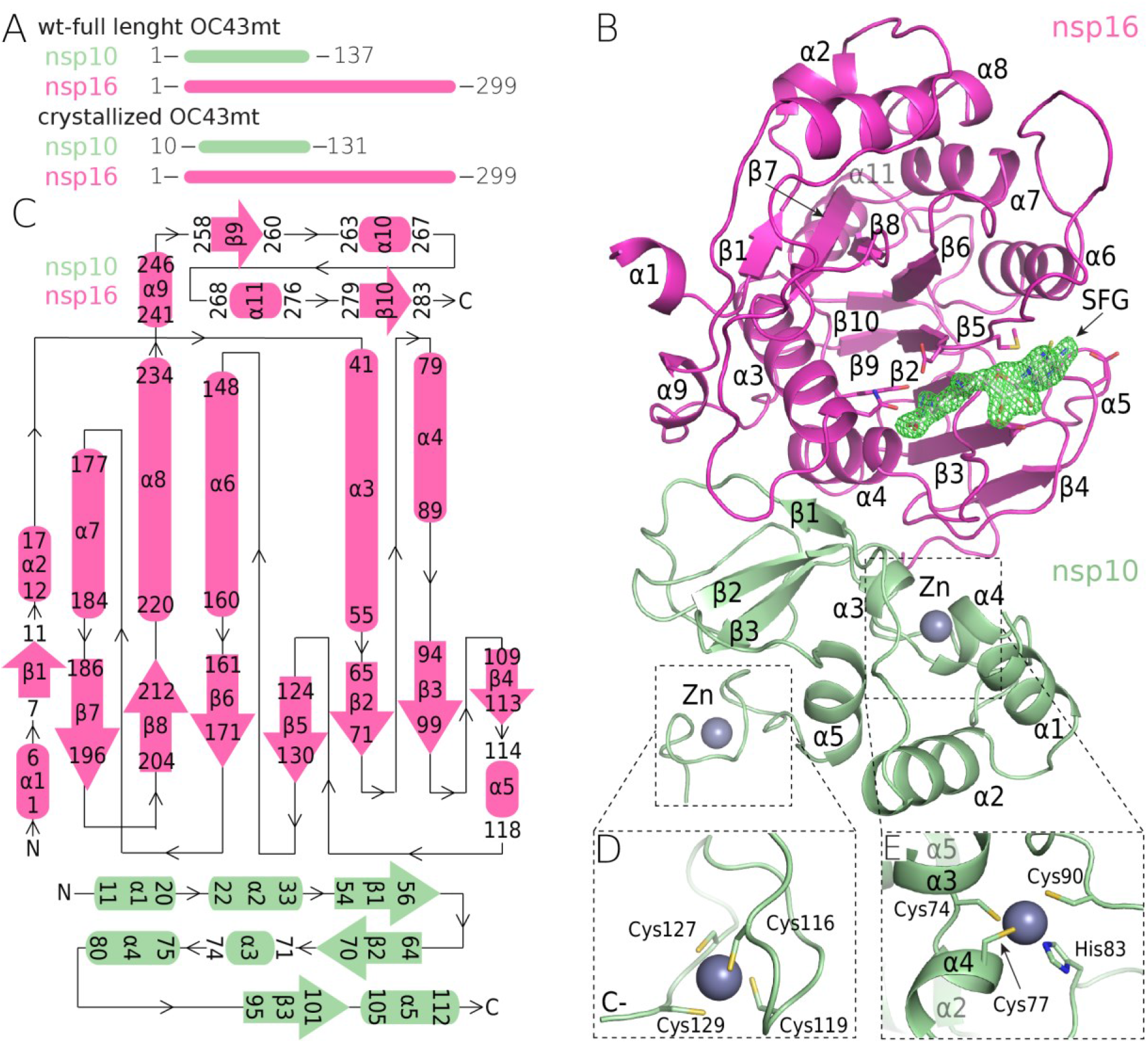
The crystal structure of the methyltransferase complex (nsp10:nsp16) from OC43-CoV. A) Schematic representation of the crystallized and full-length nsp10-nsp16 proteins; B) the overall crystal structure of nsp10-nsp16 complex (shown as ribbon) with a sinefungin molecule (white) bound in the active site (SFG), a “kick-out” omit map (Fo-Fc) of electron density (green) at 3.5 sigma with SFG excluded from the calculation; C) a topology representation of the secondary structure features of the complex; and D-E) the Zn^2+^ coordination centers in nsp10.

### SAM binding pocket

Prior to crystallization trials the nsp10:nsp16 2’-O MTase was supplemented with 1 mM sinefungin, an adenosine derivative originally isolated from *Streptomyces griseoleus* by Eli Lilly and Co. as a potential antibiotic [20] that is a pan-methyltransferase inhibitor. The electron density for sinefungin is clearly visible (Figure 2B). Detailed analysis revealed residues that are responsible for sinefungin binding. Asp114 binds the adenine aminogroup while Asp99 forms hydrogen bonds with both ribosyl hydroxyl groups. The aminoacid moiety of sinefungin is coordinated by hydrogen bonds to Asp130 and Tyr47 while Asn43 and His69 contribute to ligand binding via water bridges (Figure 3). All of these residues are absolutely conserved in human coronaviruses (Figure 1) even the Asn43 and His69 that contribute to ligand binding indirectly.

**Figure 3.**
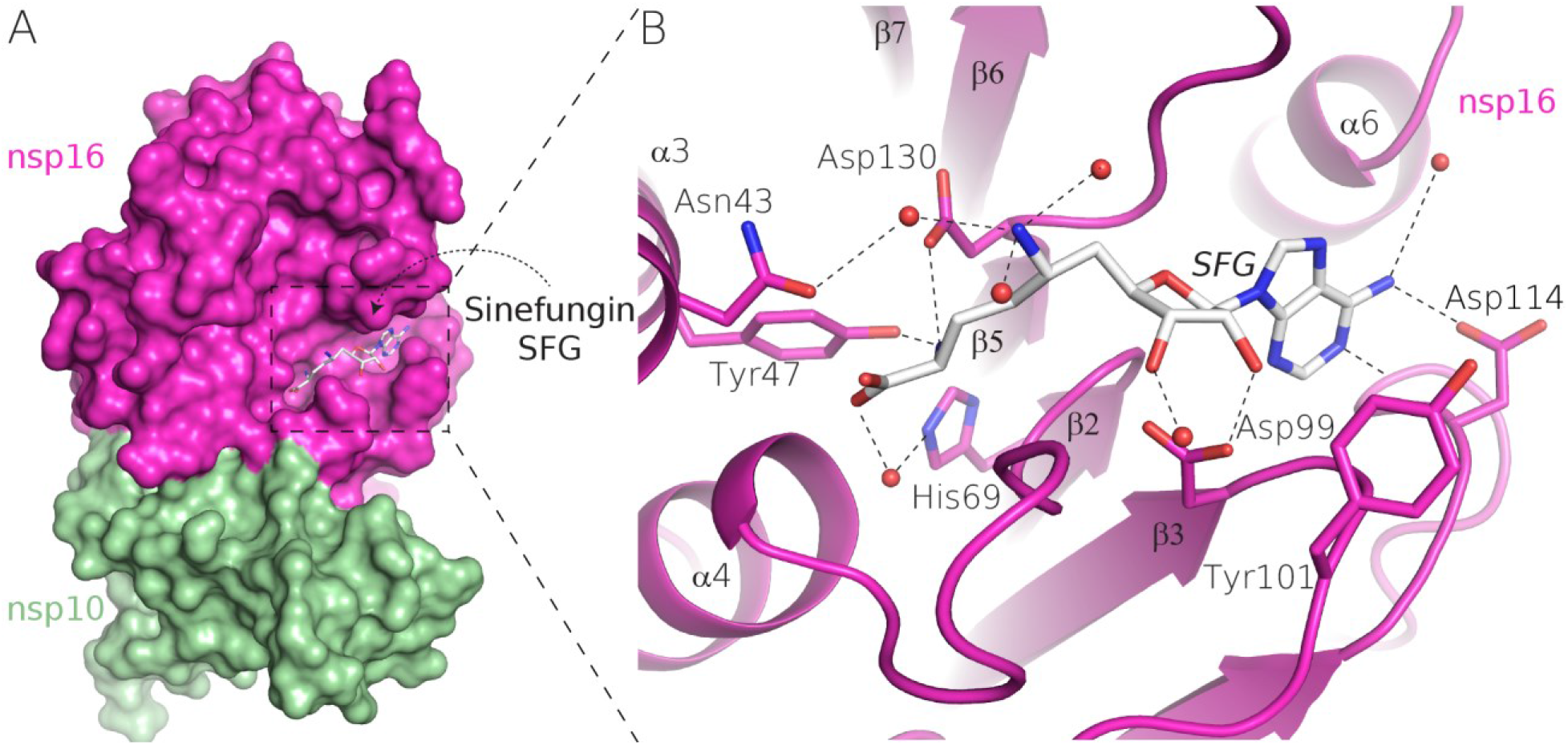
Detail of sinefungin binding to nsp16. A) the overall surface representation of the nsp10:nsp16 complex of OC43CoV, B) a detailed view of the SFG interaction with the active site of the enzyme. The amino acids involved in the interaction are shown as sticks, the waters are shown as red spheres, and the hydrogen bonds are depicted as dashed lines.

### Comparison to SARS-CoV-2 nap10:nsp16 MTase

We compared the structures of OC43 and SARS-CoV-2 nsp10:nsp16 MTases to gain insight into structural conservation of this enzyme within human coronaviruses. The overall fold remains conserved with RMSD 0.99 Å^2^ for the nsp16 and 1.03 Å^2^ for the nsp10 proteins (Figure 4A) as is the sinefungin binding pocket with the exception of Tyr101 where most human CoVs (SARS-, SARS2-, MERS-, NL63-, and HKU1-CoV) have an Asn residue instead (Figure 1). Interestingly, the position of several water molecules within the SAM binding pocket is also conserved, suggesting importance of the water bridges for ligand binding (Figure 4B).

**Figure 4.**
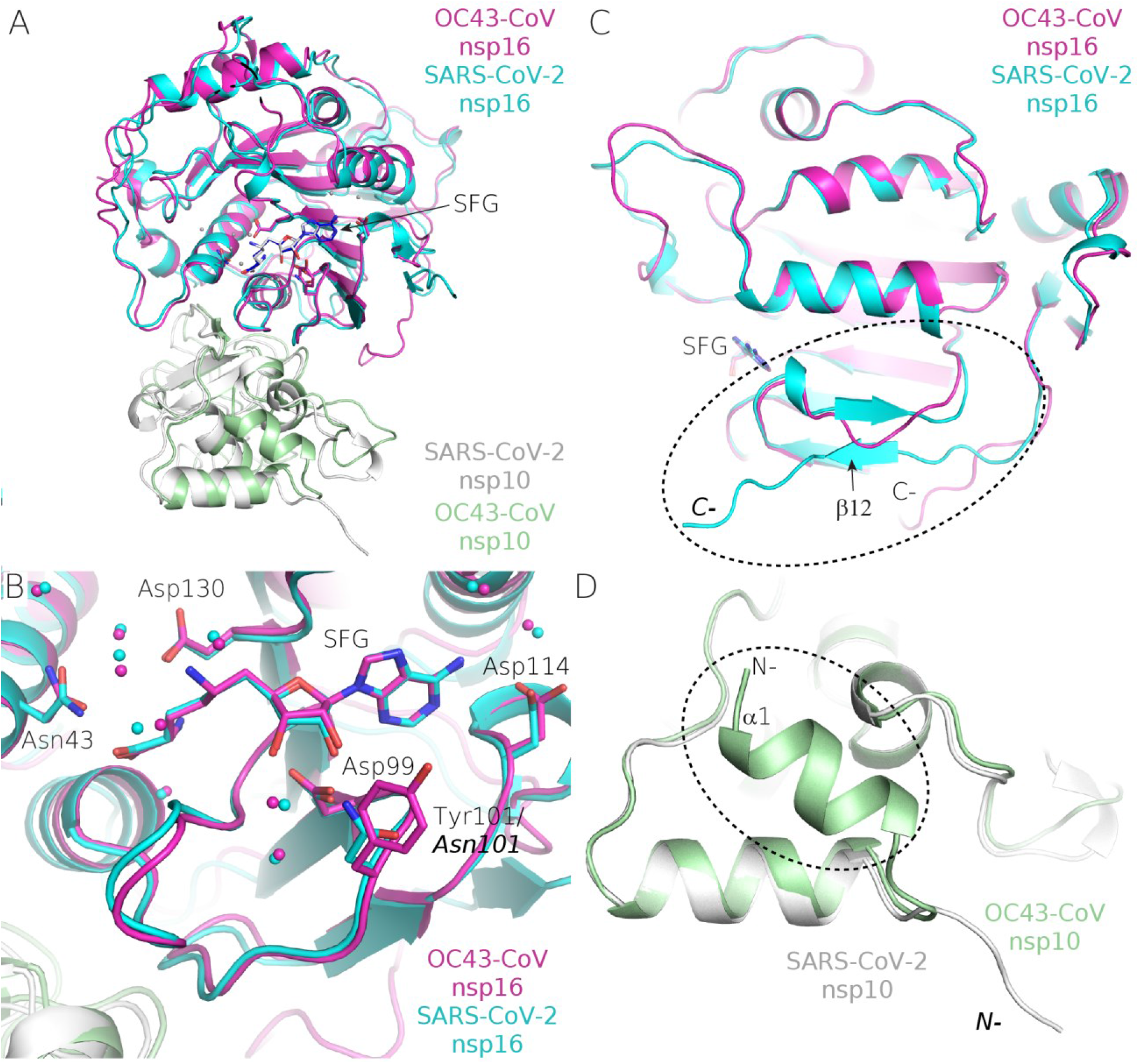
Structural alignment of the OC43-CoV and SARS-CoV2 2’-O-RNA MTase. A-B) The overall fold and orchestration of the catalytic center of the aligned SARS-Cov2 and OC43-CoV MTases and a more detailed view of the SAM binding pocket of these two protein complexes; C) the C-terminus of nsp16 of OC43- CoV (magenta) lacks the final β-sheet of nsp16 SARS-CoV2 (cyan) and it is driven toward nsp10, where it is a part of the nsp10:nsp16 interface, whilst the SARS-CoV2 C-terminus of nsp16 is involved in crystal contact (SI Figure 1) D) the N-terminal region of nsp10 of OC43- CoV (green) contains an additional α-helix. The SARS-CoV2 N-terminus is yet again involved in crystal contact (SI Figure 1).

However, the structural alignment also revealed significant differences. The C-terminus of OC43-CoV nsp16 is located at the nsp10:nsp16 interface and contributes to nsp10:nsp16 binding. While the SARS-CoV-2 nsp16 C-terminus is folded away from the interface and actually forms crystal contacts (SI Figure 1); it also has a final β-sheet (β-12) that is antiparallel to β5. Surprisingly, this β-sheet (β-12) is missing on the OC43 nsp16. On the other hand, the OC43 nsp10 has an additional α-helix (residues 11 - 20) that is not present in the SARS-CoV-2 nsp10 (Figure 4D).

## Discussion

Before the appearance of SARS and MERS, the coronaviruses were rather an insignificant group that did not attract that much attention [21], despite some of the most important discoveries, such as proteolytical processing of the Spike protein (at those times called the E2 glycoprotein) or that the leader sequence of subgenomic RNAs comes from the 5’ end of the genome, had already been made [22,23]. Upon the appearance of MERS and SARS, the coronaviruses attracted significant scientific interest that led to the molecular description of their life cycle (reviewed in Snijder et al.) [9] including structural understanding of their key enzymes [19,24–30]. However, the four human coronaviruses that generally do not cause any serious illness (OC43-, 229E-, NL63-, and HKU1-CoVs) are still understudied. Nonetheless, it would be interesting and important to understand why certain coronaviruses cause deadly diseases and other just mild colds. Certainly, the spike protein is the primary determinant of pathogenicity: both the SARS-CoV and the SARS-CoV-2 spike proteins recognize angiotensin-converting enzyme 2 (ACE2) as the receptor [31,32], while the MERS-CoV uses dipeptidyl peptidase 4 [33], the OC43- and the HKU1-CoVs use 9-*O*-acetylated sialic acids [34] and E229-CoV uses aminopeptidase N [35]. However, NL63-CoV also uses ACE2 as its receptor [36,37] and causes only common colds, suggesting that other determinants of coronaviral perilousness also exist.

Here, we reported the structure of the human OC43-CoV 2’-O MTase, a complex of two non-structural proteins: nsp10 and nsp16. The main function of this complex is to create the cap structure on the 5’ end of viral RNA. The structure revealed that despite significant sequence differences between OC43 and other human coronaviruses, the overall fold is well preserved: the biggest difference is an additional helix (α1) in the nsp10 subunit that is not observed in SARS-CoV and SARS-CoV-2 coronavirus and also a different conformation of the OC43 nsp16 C-terminus. The C-terminus is also missing a β-sheet compared to its SARS-CoV-2 counterpart. Importantly, the structure revealed high conservation of the SAM binding pocket. All residues that make direct contact with the ligand (sinefungin) are absolutely conserved among human coronaviruses suggesting that mutations of these residues are not tolerated. This is definitely good news for drug design because compounds that target the SAM pocket will likely be effective against all human coronaviruses and the evolution of resistance against them is unlikely.

## Materials and methods

### Protein expression and purification

The genes encoding for OC43 nsp10 and nsp16 proteins were codon optimized for *E. coli* and commercially synthesized (Thermo Fisher Scientific). The genes were cloned in a home-made pSUMO-Kan plasmid (a pRSFD derived vector already encoding a His_8x_-SUMO solubility and purification tag). The proteins were purified using our established protocols for viral enzymes [38,39]. Briefly, *E. coli* BL21 cells were transformed with nps10 and nsp16 expression vectors and cultivated at 37 °C in LB medium with addition of ZnSO_4_ (10 μM) and kanamycin (50 μg/ml). The production of nsp16 and nsp10 proteins was induced by addition of IPTG (300 μM) at the late exponential growth phase (OD = 0.6-0.8) and the temperature was lowered to 18 °C for 18 hours. The bacterial cells were collected by centrifugation, resuspended in lysis buffer (50 mM Tris pH 8, 300 mM NaCl, 20 mM Imidazole; 5 mM MgSO_4_; 3 mM β - mercaptoethanol; 10% glycerol) and lysed by sonication. The lysate was centrifuged and the supernatant was incubated with Ni^2+^ agarose (Macherey - Nagel), washed with lysis buffer and the recombinant proteins were eluted by the lysis buffer supplemented with 300mM imidazole. The His_8x_-SUMO tag was cleaved by the Ulp1 protease during dialysis against the lysis buffer. After dialysis the His_8x_-SUMO tag was removed by a second round of affinity chromatography. The proteins were further purified by gel filtration on a Hiload 16/600 Superdex 75 column (GE Healthcare) in the size exclusion buffer (50 mM Tris pH 7.4, 150 mM NaCl, 1 mM TCEP, 5% glycerol). The purified proteins were mixed in 1:1 molar ration, concentrated to 5 mg/ml and immediately used for crystallographic trials. After purification proteins were concentrated to 5 mg/ml and stored at −80 °C.

### Crystallization and structural analysis

The nsp10:16 protein complex was supplemented by 1 mM sinefungin Crystals of the nsp10:nsp16 protein complex grew in three days in commercial JCSG I-IV screens (Qiagen). They were cryo-protected in mother liquor supplemented with 20% glycerol and flash frozen in liquid nitrogen. The crystals diffracted to 2.2Å resolution and belonged to the trigonal P31 spacegroup. Data were collected using a home source (Rigaku) equipped with a Pilatus detector (Dectris). The data were indexed, scaled and integrated using XDS [40]. The structure was solved by molecular replacement using the structure of the SARS-CoV-2 nsp10:nsp16 (PDB id 6YZ1) as the search model in Phaser [41] and refined in Phenix [42] together with manual building in Coot [43]. The structure was refined to good R factors (R_work_ = 16.41%, R_free_ = 21.01%) and good geometry as summarized in Table 1. RMSDs were also calculated in Coot using the secondary structure matching algorithm.

## Acknowledgment

The project was supported by the Czech Science Foundation (grant number: 21-25280S), the support of the Academy of Sciences of the Czech Republic (RVO: 61388963) is also acknowledged. We are grateful to Mike Downey for critical reading of the manuscript.

## Author contribution

P.D., P.K. and E.B. performed experiments, J.S. and E.B. analyzed data, E.B. designed and supervised the project and wrote the manuscript.

